# Induction of human trophoblast stem-like cells from primed pluripotent stem cells

**DOI:** 10.1101/2021.07.14.452371

**Authors:** Yu Jin Jang, Mijeong Kim, Bum-Kyu Lee, Jonghwan Kim

## Abstract

The placenta is a transient but important multifunctional organ crucial for healthy pregnancy for both mother and fetus. Nevertheless, limited access to human placenta samples and the paucity of a proper *in vitro* model system has hampered our understanding of the mechanisms underlying early human placental development and placenta-associated pregnancy complications. To overcome these constraints, we established a simple procedure with a short-term treatment of bone morphogenetic protein 4 (BMP4) in trophoblast stem cell culture medium (TSCM) to convert human primed pluripotent stem cells (PSCs) to trophoblast stem-like cells (TSLCs). These TSLCs show not only comparable morphology and global gene expression profiles to *bona fide* human trophoblast stem cells (TSCs) but also long-term self-renewal capacity with bipotency that allows the cells to differentiate into functional extravillous trophoblasts (EVT) and syncytiotrophoblasts (ST). These indicate that TSLCs are equivalent to genuine human TSCs. Our data suggest a straightforward approach to make human TSCs directly from pre-existing primed PSCs and provide a valuable opportunity to study human placenta development and pathology even from patients with placenta-related diseases.

**HIGHLIGHTS:** Short-term treatment of BMP4 in TSCM induces human primed PSCs into TSLCs

TSLCs possess similar self-renewal and bipotency as bona fide TSCs

Global gene expression profiling shows high similarity between TSLCs and TSCs

TSLC-derived EVT and ST possess characteristics shown in TSC-derived counterparts

## 1. INTRODUCTION

The placenta is a temporary but pivotal organ that supports the fetus’s growth during pregnancy by transporting nutrients and hormones, exchanging respiratory gases and waste, and conferring immunological protection (Lewis et al., 2012). The human placenta originates from the trophectoderm, the outer layer of blastocyst, which develops into various specialized cell types including cytotrophoblasts (CT), syncytiotrophoblasts (ST), and extravillous trophoblasts (EVT) (Guttmacher et al., 2014). Recently established trophoblast stem cells (TSCs) possess an ability to differentiate into multiple cell types, such as multinucleated ST that are critical for gas/nutrient exchange as well as for metabolic and immunological functions and invasive EVT that migrate into the maternal uterus and modify its vessels to establish maternal blood flow into the placenta (Okae et al., 2018). Abnormal differentiation of trophoblasts and dysregulation in placental functions endangers the mother and fetus, leading to diverse pregnancy complications such as preeclampsia, stillbirth, miscarriage, and intrauterine growth restriction (Fisher, 2004). Notably, these disorders not only increase infant mortality and maternal morbidity but also impair the lifelong health of newborns (Lacroix et al., 2013). However, practical and ethical restrictions on accessing human placenta samples during pregnancy have greatly hindered precise understanding of the mechanisms underlying placenta development and placenta-associated complications.

Due to such limitations, animal models and *in vitro* animal model cell lines have been extensively used in the field. However, to fully understand the mechanisms underlying human placentation and placenta-related disorders, it is necessary to establish a reliable *in vitro* human model system that can mimic human placenta development. Although human TSCs have been successfully generated from both the blastocysts and CT from the first-trimester of placenta (Okae et al., 2018), from a clinical perspective, sourcing TSCs would not be straightforward due to ethical issues and health risks to both the mother and fetus when explanting placenta samples. As an alternative source of human TSCs, several attempts have been made to convert human PSCs including embryonic stem cells (ESCs) and induced pluripotent stem cells (iPSCs) to trophoblast lineage cells using bone morphogenetic protein 4 (BMP4) or BMP4-containing culture conditions (Amita et al., 2013; Horii et al., 2016; Xu et al., 2002). Although BMP4 treatment facilitate differentiation of human ESCs to trophoblasts (Li and Parast, 2014; Xu et al., 2002), they often resulted in extraneous expression of mesoderm or amnion markers (Bernardo et al., 2011; Guo et al., 2021) and a failure of establishing long-term maintainable self-renewing bipotent cells. Upon establishment of *bona fide* human TSC culture conditions, there were several attempts to generate self-renewing human TSC-like cells (TSLCs) from PSCs. A recent paper reported that human iPSCs can generate TSLCs under a micromesh culture condition (Li et al., 2019). However, this method is time-intensive and still resulted in a heterogeneous population of ST and EVT after differentiation. Another study reported the generation of two distinct trophectoderm lineage stem cells (CDX2^+^ and CDX2^-^) from human PSCs in chemically defined culture conditions (Mischler et al., 2021). Notably, recent studies have reported that TSLCs can be generated from human naïve PSCs or expanded potential stem cells (EPSCs) (Dong et al., 2020; Gao et al., 2019). So far, it is unclear which protocol is better to generate functional TSLCs comparable to *bona fide* TSCs. In this study, we successfully convert human primed PSCs to bipotent, self-renewing TSLCs that can differentiate into EVT and ST using trophoblast stem cell culture medium (TSCM) together with transient treatment with BMP4. The resultant TSLC-derived EVT and ST express the representative marker genes, hormones, and transporters according to their respective lineages. Global transcriptome analysis and functional assays confirm high similarity between TSLCs and TSCs. In sum, we report a simple, time- and cost-efficient method to directly convert human primed PSCs to TSLCs that are functionally equivalent to *bona fide* TSCs.

## 2. MATERIALS AND METHODS

### 2.1. Cell culture

Human PSCs (Yamanaka retrovirus reprogrammed human iPSCs, ACS-1023, ATCC and human ESCs, H9, WiCell) were maintained on Matrigel (corning)-coated plates with mTeSR1 medium (Stemcell Technologies). When the cells reached about 70∼80% confluence, they were passaged with ReLeSR (Stemcell Technologies) according to the manufacturer’s protocol, at a 1:10 ratio. Human TSCs obtained from Dr. Takahiro Arima (CT27 derived from CT isolated from the first trimester human placenta) were cultured as previously described (Okae et al., 2018). Briefly, the TSCs were plated on 5 µg/ml Collagen IV (corning)-coated dishes with TSCM (DMEM/F12 (Gibco) medium supplemented with 1% ITS-X supplement (Gibco), 0.5% Penicillin-Streptomycin (Gibco), 0.3% BSA (Sigma-Aldrich), 0.2% FBS (GeminiBio), 0.1 mM β-mercaptoethanol (Sigma-Aldrich), 0.5 µM A83-01 (Wako Pure Chemical Corporation), 0.5 µM CHIR99021 (Selleck Chemicals), 0.5 µM SB431542 (Stemcell Technologies), 5 µM Y27632 (ROCK inhibitor, Selleck Chemicals), 0.8 mM VPA (Wako Pure Chemical Corporation), 50 ng/ml EGF (PeproTech), and 1.5 µg/ml L-ascorbic acid (Sigma-Aldrich)). When the cells were 70∼80% confluent, they were detached with TrypLE (Gibco) for 10∼15 mins at 37°C and then the dissociated cells were plated onto Collagen IV-coated dishes at 1:3∼1:5 split ratio. The culture medium was changed every 2 days. All cells were incubated at 37°C and 5% CO2. For continuous BMP4 or BAP treatment (**Supplemental Figure 1**), PSCs were cultured as previously reported (Xu et al., 2002; Yabe et al., 2016). Briefly, PSCs were passaged with ReLeSR and plated on Matrigel-coated plates with mTeSR1 medium. After day 1, the medium was replaced with 100 ng/ml of human BMP4 (Gibco)-containing MEF-CM (Mouse Embryonic Fibroblast-Conditioned Medium) or BAP (10 ng/ml BMP4, 1 µM A83-01, and 0.1 µM PD173074) in DMEM/F12 supplemented with 20% knockout serum replacement (SR, Gibco) and cultured for up to 14 days. Media was replaced every day.

### 2.2. Derivation of stable TSLCs from PSCs

PSCs were detached using TrypLE and seeded on Matrigel-coated plate (3∼4×10^4^ cells/cm^2^) containing mTeSR1 medium supplemented with 10 µM Y27632. One day after seeding, the cells were cultured in TSCM containing 10 ng/ml BMP4 for 2 days (for TSLCs) or indicated days as shown in **Figure 1A**, then they were maintained on Matrigel-coated plates containing TSCM in the absence of BMP4. During the treatment of BMP4, media was changed daily. The cells were passaged every 2∼3 days when the confluency reached up to 80-90% and analyzed at day 14 or later. All cells were incubated at 37°C in 5% CO2.

**Figure 1.**
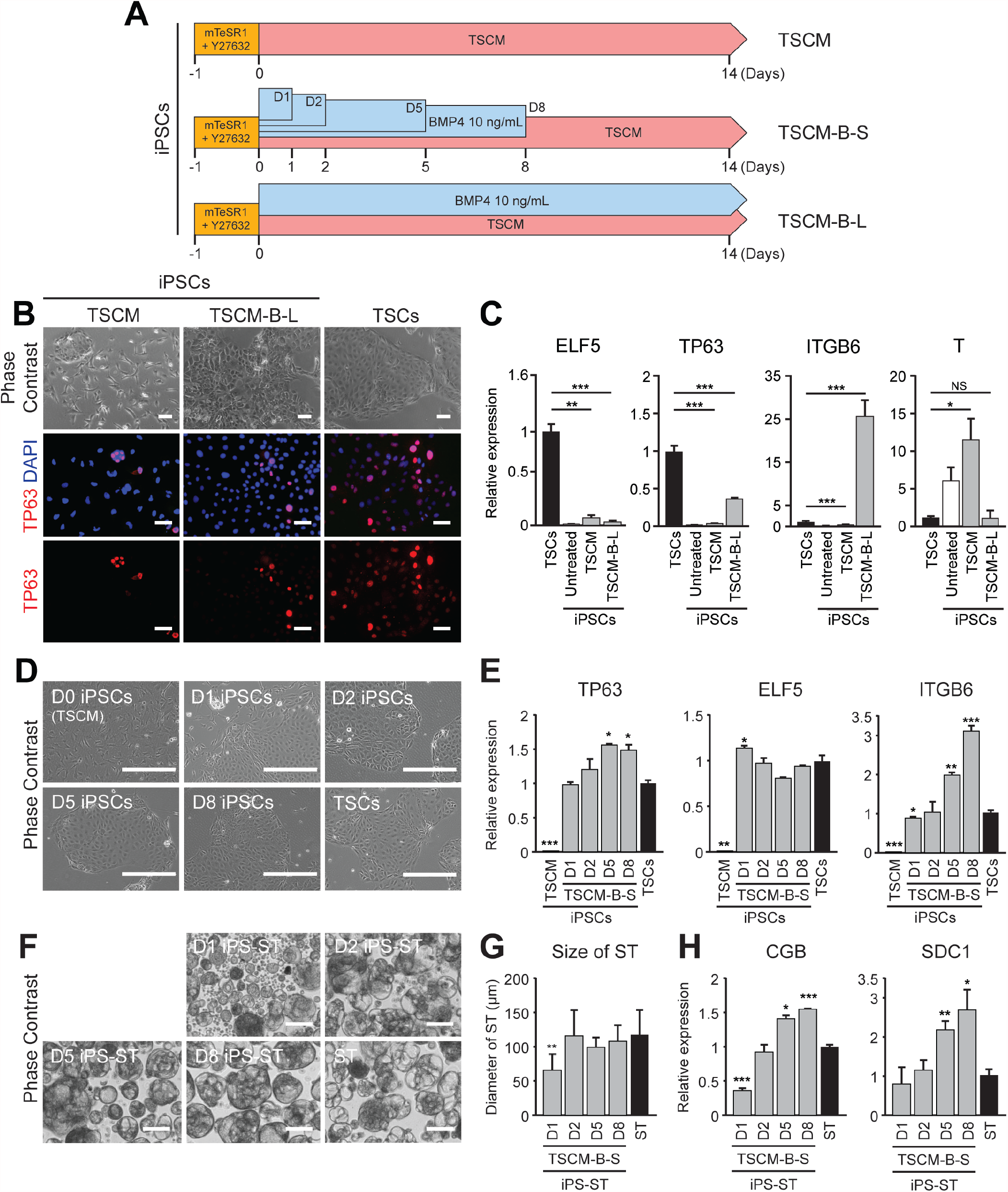
Two days of BMP4 treatment with long-term culture in TSCM is sufficient to establish TSLCs from PSCs. A) Schematic diagram of induction protocol. TSCM is used as a basal medium. There are three conditions tested to induce TSCs from human iPSCs. One is to culture the iPSCs in TSCM for over 14 days (TSCM), another is to culture the iPSCs in TSCM with BMP4 10 ng/ml for the first 1, 2, 5, or 8 days and replace TSCM without BMP4 (TSCM-B-S; D1, D2, D5, and D8) and the other is to culture the iPSCs in TSCM with BMP4 10 ng/ml for the entire duration (TSCM-B-L). B) Representative phase contrast (upper panel) and IF images (middle and bottom panels) of iPSCs cultured in TSCM or TSCM-B-L conditions and TSCs. Red indicates the protein expression of TP63. Blue (DAPI) indicates the nuclei. Overlayed images are shown in the middle panel. Scale bars indicate 100 µm. C) Relative transcript levels of TSCs (ELF5 and TP63), amnion (ITGB6), and mesendoderm (T) marker genes in the indicated cells. Error bars indicate SEM (n=3). Significance was indicated with *: p < 0.05, **: p < 0.01 and ***: p < 0.001. D) Representative phase contrast images of iPSCs cultured in TSCM or TSCM-B-S conditions (D1, D2, D5, and D8). TSCs were showed as a control. Scale bars indicate 400 µm. E) Relative transcript levels of TSCs (TP63 and ELF5) and amnion (ITGB6) marker genes in the indicated conditions. Error bars indicate SEM (n=2). Significance was indicated with *: p < 0.05, **: p < 0.01 and ***: p < 0.001. F) Representative phase contrast images of iPS-ST from the indicated conditions. The ST were differentiated from TSCs. Scale bars indicate 100 µm. G) Bar graph showing diameter of 3D-cultured iPS-ST and ST from each condition. Significance was indicated with **: p-value < 0.01. H) Relative transcript levels of ST marker genes (CGB and SDC1) in the indicated conditions. Error bars indicate SEM (n=2). Significance was indicated with *: p < 0.05, **: p < 0.01 and ***: p < 0.001.

### 2.3. EVT and ST differentiation

TSCs and TSLCs were differentiated into EVT and ST as described previously (Okae et al., 2018). For differentiation of ST, dissociated TSCs and TSLCs were cultured in a 60 mm petri dish (5×10^5^ ∼1×10^6^ cells/dish) with 4 ml ST differentiation medium (DMEM/F12 medium supplemented with 1% ITS-X supplement, 0.5% Penicillin-Streptomycin, 0.3% BSA, 0.1 mM β-mercaptoethanol, 2.5 µM Y27632, 2 µM forskolin (Selleck Chemicals), and 4% SR). ST differentiation medium was replaced on day 3. ST derived from TSCs and TSLCs were analyzed on day 6. For differentiation of EVT, dissociated TSCs and TSLCs were plated on 1 µg/ml Collagen IV-coated 6 well plate (1∼1.5×10^4^ cells/cm^2^) with 2 ml of EVT differentiation medium (DMEM/F12 medium supplemented with 1% ITS-X supplement, 0.5% Penicillin-Streptomycin, 0.3% BSA, 0.1 mM β-mercaptoethanol, 100 ng/ml human neuregulin-1 (NRG1, Cell Signaling Technology), 7.5 µM A83-01, 2.5 µM Y27632, 4% SR, and 2% Matrigel). On day 3, the EVT medium was replaced with fresh medium supplemented with reduced Matrigel (0.5%) in the absence of NRG1. On day 6, the EVT medium was replaced again with the medium containing reduced Matrigel (0.5%) but omitting both NRG1 and SR. EVT derived from TSCs and TSLCs were analyzed on day 8.

### 2.4. RNA extraction and RT-qPCR assay

Total RNA was extracted using the RNeasy plus mini kit (Qiagen) following the manufacturer’s instructions and cDNAs were synthesized from 300-500 ng of RNAs by qScript™ cDNA SuperMix (Quantabio). RT-qPCR was performed with PerfeCTa® SYBR® Green FastMix®, Low ROX™ (Quantabio) and 2 µl of 15-20X diluted cDNA. RT-qPCR primer sequences are listed in **Table S1**. The results were analyzed by ΔΔCt method to calculate fold changes with normalization against GAPDH.

### 2.5. RNA sequencing (RNA-seq)

To quantify global gene expression, RNA-seq was performed using PSCs, TSLCs, TSCs and their derivatives. Total RNAs were extracted using the RNeasy plus mini kit following the manufacturer’s instructions. RNA-seq libraries were generated with 300 ng of total RNA using NEB Next Ultra II RNA Library prep kit (NEB, E7770) according to the vendor’s protocol. Briefly, mRNAs were isolated from total RNAs with Magnetic mRNA Isolation Kit (oligo(dT) beads) (NEB, E7490). First strand cDNAs were synthesized using random primers and second strand cDNAs were synthesized subsequently. Double stranded cDNAs were purified by NEBNext Sample Purification Beads (NEB, E7767), and then the end prep reaction was performed followed by ligation with sample-specific adaptor and index. RNA-seq libraries were sequenced using an Illumina Novaseq 6000 machine. Paired-end reads from RNA-seq were aligned to the reference human genome (hg38) using STAR (v2.5.2b). To generate read counts for each gene, the python package HTSeq (https://htseq.readthedocs.io/en/master/) was used. Counts were quantified by the R package DESeq2 (v1.28.0) (Love et al., 2014) and regularized log transformed using the rlog function. Heat maps were generated using Java TreeView (v1.1.6r4).

### 2.6. RT-qPCR for miRNAs

miRNAs and total RNAs were isolated using the miRNeasy mini kit (Qiagen) according to the manufacturer’s instructions. 1 µg of RNAs was converted to cDNAs by using TaqMan™ MicroRNA Reverse Transcription Kit (Thermo Fisher Scientific). RT-qPCR was conducted with PerfeCTa® SYBR® Green FastMix®, Low ROX™. miRNA primer sequences for RT-qPCR are listed in **Table S1**. The data were analyzed by ΔΔCt method to calculate fold changes with a normalization against miR-103a.

### 2.7. Growth curve

The TSLCs and TSCs were plated on 96-well plates. Cell Counting Kit-8 (Dojindo Laboratory) was used for a cell proliferation assay. Every 24 hr, 10 µl of Cell Counting Kit-8 solution was added to the cultured medium and incubated for 2 hrs. The absorbance was measured at 450 nm with Infinite® M1000 PRO microplate reader (Tecan) for 6 days.

### 2.8. ELISA assay

The cultured media was collected from ST derived from TSLCs and TSCs on differentiation day 6. Secretion of human chorionic gonadotropin (hCG) was analyzed by enzyme-linked immunosorbent assay (ELISA) kit for hCG (RayBiotech) according to the manufacturer’s instructions.

### 2.9. Immunofluorescent (IF) staining

The cells were cultured in a µ-Slide 8 Well chamber (ibidi) for 2 days (TSLCs and TSCs) or 8 days (EVT), and then fixed with 4% paraformaldehyde (PFA; Sigma-Aldrich) for 20 mins at room temperature (RT). The ST were cultured for 6 days in petri dish and collected into 1.5 ml tube. The ST were fixed with 4% PFA for 1 hr at RT. The fixed cells were permeabilized and blocked with 0.3% Triton X-100 (Sigma-Aldrich) and 10% normal goat serum (Sigma-Aldrich) in Dulbecco’s phosphate-buffered saline (PBS) (Thermo Fisher Scientific) for 45 mins at RT. For HLA-G staining, a blocking solution was used without 0.3% Triton X-100. The samples were incubated with diluted primary antibodies (**Table S2**) at 4°C for overnight. The cells were incubated with Alexa Fluor 488- or Alexa Fluor 594-conjugated goat secondary antibodies (1:400; Invitrogen) for 1 hr 30 mins at RT. The nuclei of the cells were counterstained with 4’,6-diamidino-2-phenylindole (DAPI, Sigma-Aldrich) for 5 min and the images were acquired using a fluorescence microscope (Zeiss Axiovert, Carl Zeiss).

### 2.10. Bisulfite sequencing

For extraction of genomic DNA, the cells were resuspended with Tris-HCl (pH 8.0; Sigma-Aldrich), 10 mM EDTA (Thermo Fisher Scientific), 0.5% SDS (Thermo Fisher Scientific), and 200 µg/ml proteinase K (New England Biolabs) and incubated for 2 hrs at 55°C. 0.2 M NaCl was added to the tube. The cell lysates were mixed with PCI (Phenol:Chloroform:Isoamyl Alcohol, Invitrogen) by following manufacturer’s instruction and centrifuged with Phase Lock gel (QuantaBio) for 2 mins at 12,000 rpm. The supernatant was moved into a new tube. 25 µg/ml ribonuclease A (Life Technologies) was added, and the tube was incubated at 37°C for 1 hr. The cells lysates were mixed with PCI and centrifuged with Phase Lock gel again for 2 mins at 12,000 rpm. The supernatant was mixed with 2.5X volume of 100% ethanol for precipitation of genomic DNA and centrifuged for 5 mins at 12,000 rpm. The genomic DNA was resuspended in 10 mM Tris-HCl, 1 mM EDTA. One µg of genomic DNA was used for bisulfite conversion by using EpiTect Bisulfite Kit (Qiagen) according to the manufacturer’s instructions. Converted DNA was 10X diluted with 10 mM Tris-HCl, 1 mM EDTA. The ELF5 promoter region was amplified using primers as described previously (Lee et al., 2016). Each PCR product was amplified using EpiMark Hot Start Taq DNA Polymerase (New England Biolabs) following the manufacturer’s protocol. The PCR products were purified with a MinElute PCR purification kit (Qiagen) and cloned into qCR2.1 Vector from TA Cloning Kit (Thermo Fisher Scientific) using a molar ratio of 1 plasmid to 2 inserts. After transformation into DH5α competent bacteria, miniprepped DNAs using PureLink™ Quick Plasmid Miniprep Kit (Invitrogen) were analyzed by Sanger sequencing.

### 2.11. Invasion assay

The 3D invasion assay was performed by modifying a previously published protocol (Bayless et al., 2009). Molecular Probes Secure Seal Hybridization Chamber (Thermo Scientific) were treated with ultraviolet (UV) light for 1 hr in a laminar flow hood. The chambers adhered to the bottom of 12-well plate. Matrigel and medium (TSCM or EVT differentiation medium) mix were prepared as 2:1 ratio on ice. 20 µl of each Matrigel mix were added to one of the holes in the chamber and equilibrated for 1 hr at 37°C. 5×10^4^ cells of TSLCs and TSCs were seeded with 30 µl of TSCM or EVT differentiation medium into another hole of the chamber. In order to sink the cells to the Matrigel mix, the plate was standing on the side and incubated for 1 hr at 37°C. After then, the chamber was completely covered with 1 ml of additional culture medium. And then, the differentiation was followed as described in 2.3.

### 2.12. Sodium Fluoresceine (Na-Flu) leakage assay

For evaluation of placental barrier function, 5×10^4^ cells/cm^2^ in 0.1 ml TSCM or ST differentiation medium (detailed information in 2.1 and 2.3) were introduced onto 0.4 µm pore membrane of the apical chamber (corning, 3470). The basal chamber of the transwell was filled up with 0.6 ml of TSCM or ST differentiation medium. After day 3 and day 6 post seeding, medium of apical chamber was aspirated and transferred to a new 24-well containing 1 ml of PBS for washing. Then, 100 µl of 5 µM Na-Flu in TSCM or ST differentiation medium was added to the apical chamber. The apical chamber was transferred to a new 24-well containing 600 µl of DMEM/F12 medium (without any growth factor supplements) for 1 hr in 5% CO2 at 37°C. 50 µl of samples from the basal chamber were pipetted into a 96-well plate and analyzed using an Infinite® M1000 PRO microplate reader (Ex. 460 nm/ Em. 515 nm; Tecan). After measurement, the fluorescein solution was aspirated and replaced with TSCM or ST differentiation medium.

### 2.13. Flow cytometry

EVT cells were differentiated from iPS-TSLCs, ES-TSLCs and TSCs for 8 days. The cells were dissociated by TrypLE at 37°C for 30 mins and diluted with DMEM/F12. After washing with FACS buffer (PBS without Mg^2+^, Ca^2+^, 5% FBS, 2 mM EDTA, and 0.1% sodium azide), the cells were incubated with Phycoerythrin (PE)-conjugated HLA-G (Abcam) 1:50 diluted in FACS buffer for 30 mins on ice. After incubation, the cells were washed 3 times with FACS buffer in the dark. Flow cytometry analysis was performed using a BD Accuri (BD Biosciences), and the data were analyzed using FlowJo software (LLC).

### 2.14. Statistical analysis

Unless otherwise described, all statistical analyses were performed with three biological replicates. Bar chart results were presented as the mean with the standard error of the mean (SEM). Student’s t-test was used to determine the significance of differences between groups with the annotations: *: p < 0.05, **: p < 0.01, and ***: p < 0.001.

### 2.15. Data sets used for analysis

We downloaded expression data from GSE66302 (human amnion at 9 gestational weeks), to compare gene expression profile to TSLCs.

### 2.16. Data deposit

Raw and processed RNA-seq data have been deposited to the public server GEO database under accession number GSE178162.

## 3. RESULTS

### 3.1. Short-term treatment of BMP4 is crucial to generate homogeneous and long-term maintainable human TSLCs from primed PSCs

Multiple previous studies showed that the treatment of BMP4 or BAP (combination of BMP4, A84-01, and PD173074) can convert ESCs into trophoblast lineage cells (Amita et al., 2013; Horii et al., 2016; Xu et al., 2002; Yabe et al., 2016), indicating that BMP4 triggers trophoblast-specific gene expression programs. However, the BMP4- and BAP-treated cells were differentiated into heterogeneous cell populations which contained ST- and EVT-specific marker expressing cells and failed to acquire self-renewal capacity. In accordance with the previous observations, our continuous treatment of BMP4 or BAP to iPSCs (see Materials and Methods) resulted in mixed cell populations. In particular, the BMP4-treated cells start to die after 10 days of culture, indicating the loss of self-renewing ability (**Supplemental Figure 1A**). Although the BAP-treated cells were more proliferative than the BMP4-treated cells, they did not show a typical morphology of TSCs (**Supplemental Figure 1A**). The expression levels of TSC markers (TP63 and ELF5) were only 15-30% of the control TSCs in both BMP4- and BAP-treated cells, even though the expression levels were induced compared to undifferentiated iPSCs. (**Supplemental Figure 1B**). In addition, The BMP4-treated cells expressed a higher level of mesendoderm marker (T), while the BAP-treated cells did not upregulate the expression of T (**Supplemental Figure 1B**). Furthermore, while these cells expressed higher levels of HLA-G (EVT marker) and CGB (ST marker) than undifferentiated iPSCs as described previously (**Supplemental Figure 1B**) (Xu et al., 2002; Yabe et al., 2016), the levels were significantly lower than those observed in EVT or ST differentiated from TSCs, suggesting that BMP4- and BAP-treated cells may not be able to fully differentiate into EVT or ST. With morphological heterogeneity observed in BMP4- and BAP-treated cells, all these results indicate that continuous treatment of BMP4 or BAP is not opted to generate a self-renewing population of TSLCs. In addition, while generating both EVT and ST, guided differentiation into specific lineage toward EVT or ST has not been tested.

Since BMP4 was sufficient to trigger the loss of ESCs characteristics and induce trophoblast differentiation (Amita et al., 2013; Xu et al., 2002), and TSCM maintain the self-renewal of TSCs *in vitro* (Okae et al., 2018), we subsequently tested continuous BMP4 treatment to the primed iPSCs in TSCM (TSCM-B-L) for 14 days with a control TSCM alone condition (**Figure 1A**). Cells cultured in TSCM-B-L condition showed polygonal morphology, which is typical for endodermal cells that are relatively more proliferative than cells grown in MEF-CM (basal medium for BMP4- and BAP-treated protocols). Meanwhile, cells cultured in TSCM showed only a few colony-forming cells as previously reported (Cinkornpumin et al., 2020) (**Figure 1B**). Only a few TP63-expressing cells in both conditions by immunofluorescent staining (**Figure 1B**), and the transcript levels of ELF5 and TP63 in TSCM and TSCM-B-L conditions were lower than those in control TSCs (**Figure 1C**). Notably, T was highly expressed in the cells cultured in TSCM alone (**Figure 1C**), while ITGB6 (amnion marker) was upregulated in TSCM-B-L (**Figure 1C**). These indicate that continuous BMP4 treatment induces amnion lineage markers in PSCs rather than TSC markers as previously reported (Guo et al., 2021). We thought that long term treatment of BMP4 may lead to adverse effects on attempt to generate of TSLCs from PSCs. Since it has been reported that a transient exposure to BAP (24-36 hrs) can generate cell lines which are prone to differentiate into trophoblast lineage (Yang et al., 2015), we sought to test transient treatment of BMP4 in combination with TSCM conditions. To optimize duration of BMP4 treatment, we treated BMP4 for 1, 2, 5, and 8 days (referred to as D1, D2, D5, and D8) and grew them for up to 14 days in TSCM after withdrawal of BMP4 (TSCM-B-S) (**Figure 1A**). During BMP4 treatment, we changed media every day and split the cells when the confluency reached up to 80-90%. At 14 days, we examined the expression levels of TSCs and amnion markers using RT-qPCR. Regardless of the duration of BMP4 treatment, the resultant cells showed similar morphology and expression levels of TP63 and ELF5 with those of TSCs (**Figure 1D** and **E**). However, ITGB6 expression gradually increased with longer BMP4 treatment (**Figure 1E**), indicating that the cells treated with BMP4 for more than 2 days is deleterious for PSCs to TSLCs conversion. To verify the differentiation potential of TSLCs, we differentiated the cells into ST. The cells treated with BMP4 for 1 day did not develop cystic morphology, which is a key characteristic of ST, while cells treated with BMP4 for longer than 1 day were able to form a cystic shape during ST differentiation that was comparable to TSC-derived ST (**Figure 1F** and **G**). While the expression of ST markers CGB and SDC1 increased gradually with longer BMP4 treatment (**Figure 1H**), ST derived from TSLCs treated with BMP4 for 2 days showed similar expression levels of ST marker genes to ST from TSCs. Considering marker gene expression, self-renewal, and differentiation potential, we finally selected 2 days BMP4 treatment as the best condition. These results demonstrate that 2 days of BMP4 treatment in TSCM is pivotal to convert human primed PSCs into proliferative TSLCs.

### 3.2. TSLCs recapitulate the hallmark gene expression of *bona fide* TSCs

To further validate our protocol and the authenticity of TSLCs, we generated TSLCs from iPSC (referred to here as iPS-TSLCs) as well as ESCs (referred to here as ES-TSLCs) both of which were maintained under primed conditions, and then examined the levels of multiple marker genes of human TSCs. Both iPS- and ES-TSLCs demonstrated TSC-like morphology in shape and size (**Figure 2A**), and they showed no detectable protein level of stem cell core factor OCT4 and NANOG after the conversion, while having similar levels of TP63 and KRT7 expression as TSCs (**Figure 2B** and **Supplemental figure 2A**). Moreover, the transcript levels of TSC-specific genes (ELF5, TFAP2C, HAVCR1, KRT7, TP63, and GATA3) were significantly induced to the comparable levels of those in TSCs, whereas they showed a significant reduction of POU5F1 and NANOG expression (**Figure 2C**). We further tested the expression of TSC-specific microRNA clusters (miR-525-3p, miR-526b-3p, miR-517b-3p, and miR-517-5p) and methylation status of *ELF5* promoter region as previously suggested (Lee et al., 2016) and confirmed slightly lower but robust expression of the tested miRNAs in both iPS- and ES-TSLCs (**Figure 2D**). Similar to TSCs, we also observed that the *ELF5* promoter in TSLCs was hypomethylated (**Figure 2E**).

**Figure 2.**
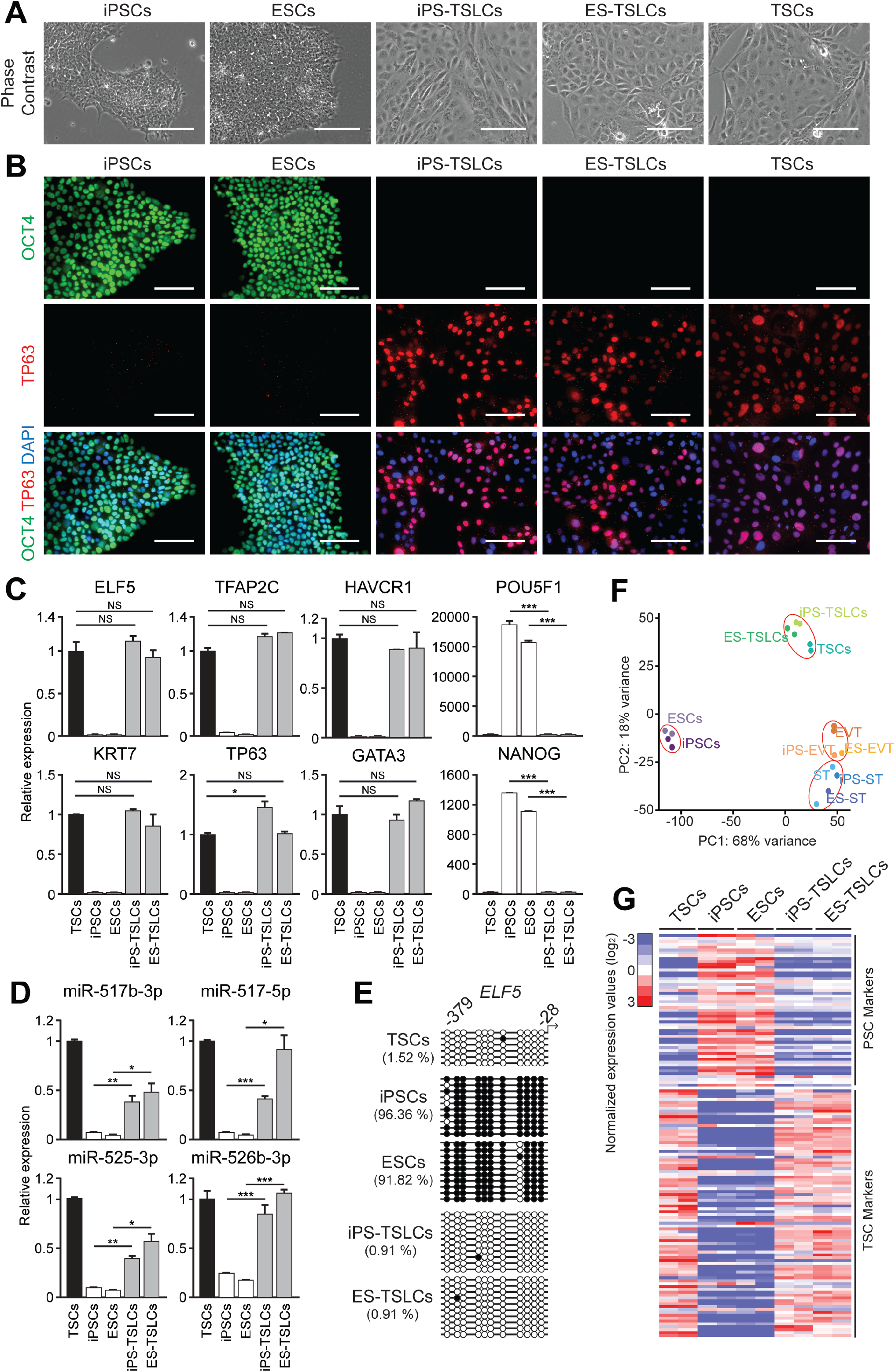
TSLCs derived from primed iPSCs and ESCs show comparable gene expression signatures to TSCs. A) Representative phase contrast images of iPSCs, ESCs (H9), iPS-TSLCs, ES-TSLCs, and control TSCs. Scale bars indicate 100 µm; all images taken at the same magnification. B) IF staining images showing the protein expression of PSC marker OCT4 (green, upper panel) and TSC marker TP63 (red, middle panel) in the indicated cell types. Overlayed images of OCT4 and TP63 are shown in the bottom panel with the nuclei staining with DAPI (blue). Scale bars indicate 100 µm. C) and D) Relative transcript levels of TSC markers (ELF5, TFAP2C, HAVCR1, KRT7, TP63, and GATA3) and PSC markers (POU5F1 and NANOG) (C) and TSC-specific miRNA clusters (miR-525-3p, miR-526b-3p, miR-517b-3p, and miR-517-5p) (D) in the indicated cell types. Error bars indicate SEM (n=3). Significance was indicated with *: p-value < 0.05, **: p-value < 0.01 and ***: p-value < 0.001. E) Distribution of methylated CpG sites (filled circles) at the *ELF5* promoter in the indicated cell types. The relative percentage of methylated CpG sites is indicated with cell types. F) Principal component analysis (PCA) for gene expression of the indicated cell types. Each dot is one biological replicate. G) A heatmap showing normalized relative expression of PSC and TSC markers (**Table S3**) in the indicated cell types indicated. Y-axis shows representative PSC and TSC markers. Each cell type displays 2 biological replicates in each column. Relative expression was calculated by dividing the level of a gene with average gene expression of all cell types.

To further evaluate whether the TSLCs possess global gene expression signatures representative of TSCs, we performed transcriptome analysis by RNA-sequencing. Principal component analysis (PCA) shows that the global gene expression profiles of iPS- and ES-TSLCs clustered clearly with *bona fide* TSCs, suggesting that TSLCs and TSCs have a similar gene expression pattern. As expected, the cluster containing TSLCs and TSCs was separated from PSCs, EVT, and ST (**Figure 2F**). Next, we confirmed that compared to PSCs, the levels of PSC-specific genes were lower in iPS-TSLCs and ES-TSLCs, while the levels of TSC-specific genes were comparable to the levels in TSCs, which further evinces proper derivation of TSLCs from PSCs (**Figure 2G, Table S3**). Since previous papers have shown that human primed PSCs differentiate into amnion-like cells upon BMP treatment (Guo et al., 2021; Zheng et al., 2019), we also monitored the levels of amnion-specific genes, which are predominantly expressed in human amnion at 9^th^ gestational weeks (Roost et al., 2015). In contrast with the amnion control, iPS- and ES-TSLCs showed similarly low expression levels of amnion genes compared to genuine TSCs (**Supplemental figure 2B, Table S3**), suggesting that our protocol did not promote PSCs to differentiate into amnion. Moreover, we confirmed that both TSLC lines can self-renew for over 20 passages without losing typical TSC morphology and proliferation capacity (**Supplemental figure 2C and D**). Consistent with the morphology, the levels of TP63, ELF5, KRT7, TFAP2C, and EGFR were stably maintained in both iPS-TSLCs and ES-TSLCs even after 20 passages (**Supplemental figure 2E**), suggesting these cells can self-renew for a long time. In summary, all these results further support that the TSLCs we generated are similar to *bona fide* TSCs.

### 3.3. TSLCs can differentiate into EVT

Since genuine TSCs can differentiate into both EVT and ST (Okae et al., 2018), we examined the differentiation potential of our TSLCs generated from primed PSCs. After EVT differentiation, both iPS-TSLC-derived EVT (iPS-EVT) and ES-TSLC-derived EVT (ES-EVT) showed primary EVT features of spindly, elongated cell morphology comparable to the control EVT derived from TSCs (**Figure 3A**). Consistently, iPS- and ES-EVT showed robust expression of EVT-specific markers of HLA-G and MMP2 in both transcript and protein levels (**Figure 3A and 3B**). We assessed the surface expression of HLA-G in EVT differentiated from iPS-TSLCs, ES-TSLCs, and TSCs using flow cytometry. Although the intensity of fluorescence signals varied between cells, 78.2% of iPS-EVT and 95.0% of ES-EVT expressed HLA-G in comparison with 91.8% of EVT differentiated from TSCs (**Figure 3C**). Next, we performed transcriptome analysis to compare EVT-specific gene expression pattern between TSLC-derived EVT and EVT. We observed that 1,123 EVT-specific genes obtained by comparing TSCs with EVT (4-fold upregulated genes) were indeed activated in TSLC-derived EVT compared to undifferentiated TSLCs (**Figure 3D, Table S3**). Since EVT invasion into the maternal decidua is a crucial step to anchor the placenta to the uterus and establish maternal blood flow into the placenta, invasion ability is a signature characteristic of EVT. To functionally validate invasion ability, we differentiated both TSLCs into EVT in Matrigel-blocked chambers (Bayless et al., 2009) (**Figure 3E**). Although the invasion efficiency of EVT from both iPS-TSLCs and ES-TSLCs seemed slightly lower than EVT derived from control TSCs, their ability to invade the Matrigel region is clear and evident compared to undifferentiated control (**Figure 3F**). These results demonstrate that primed PSC-derived TSLCs differentiate into functional EVT comparable to *bona fide* EVT.

**Figure 3.**
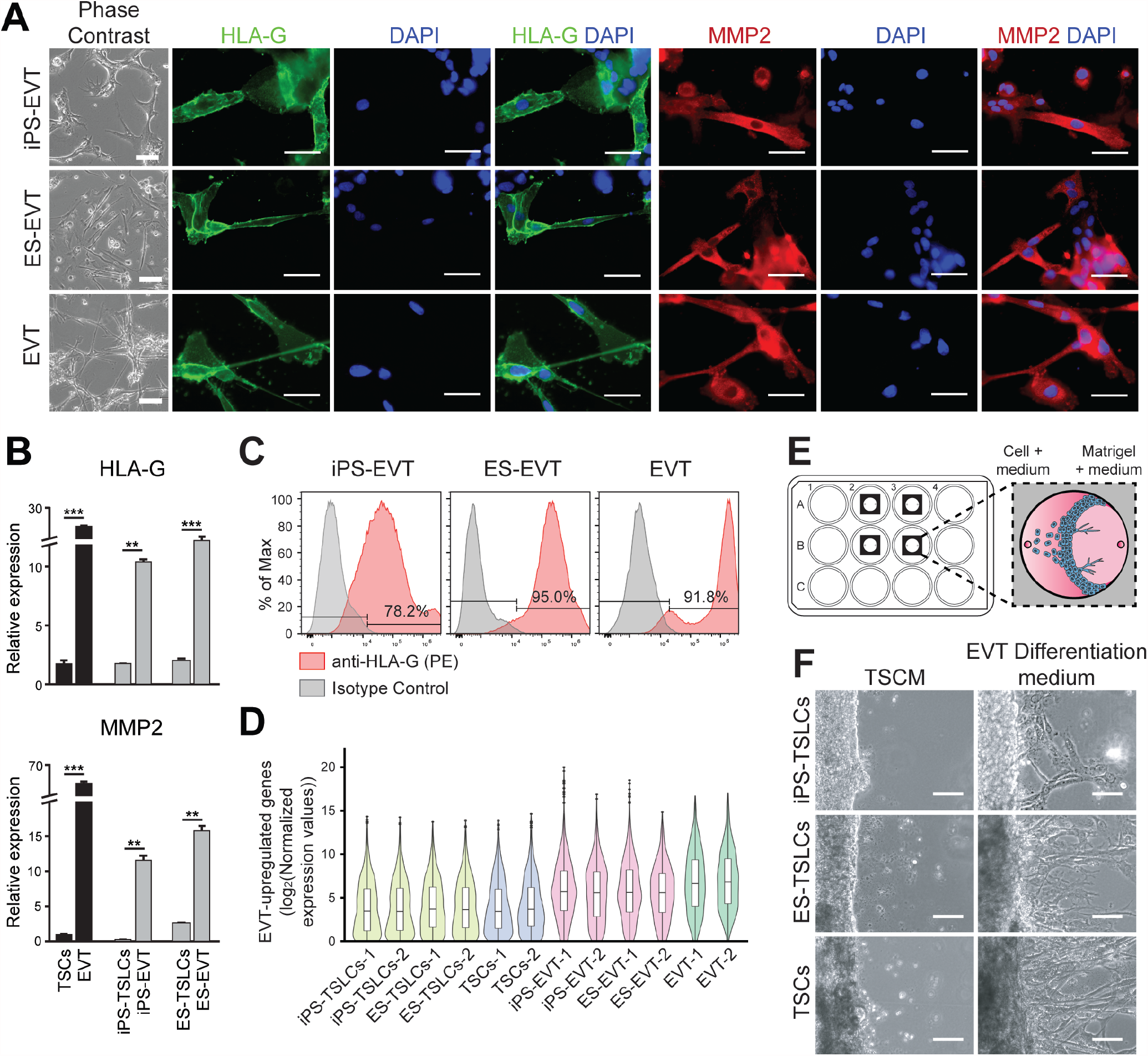
TSLCs can differentiate into EVT-like cells. A) Representative phase contrast (left panel) and IF images of indicated cell types. Green indicates the protein expression of HLA-G. Red indicates the protein expression of MMP2. Blue (DAPI) indicates nuclei. Overlayed images are shown in each right panel. Scale bars indicate 100 µm. B) Relative transcript levels of EVT marker genes (HLA-G and MMP2) in indicated cell types to control TSCs. Error bars indicate SEM (n=3). Significance was indicated with **: p < 0.01 and ***: p < 0.001. C) Flow cytometry analysis of surface HLA-G expression was performed with iPSC-EVT, ES-EVT, and EVT at day 8, which were differentiated from iPS-TSLCs, ES-TSLCs, and TSCs. The expression was compared with an isotype control of each cell type. D) Violin plots showing the expression distribution of 1,123 EVT-upregulated genes (**Table S3**) in the indicated cell types. The box indicates 25th, 50th, and 75th percentiles. E) Schematic illustration of imaging chamber. Imaging chambers were adhered to the bottom of a 12-well plate. The Matrigel and medium (TSCM or EVT differentiation medium) mix was added to one of the open ports and allowed to equilibrate at 37 °C. After 1 hr, the TSLCs or TSCs were seeded into another port with 30 µl of TSCM or EVT differentiation medium. 1 ml of TSCM or EVT differentiation medium was added to each well. F) Representative phase contrast images of invading cells in TSCM or EVT differentiation medium at day 8. Scale bars indicate 100 µm.

### 3.4. TSLCs can differentiate into ST

We also tested a differentiation potential of TSLCs toward ST. After ST differentiation, both iPS-TSLC-derived ST (iPS-ST) and ES-TSLC-derived ST (ES-ST) exhibited a cystic morphology that is a key characteristic of ST grown in 3D culture in conjunction with a robust level of CGB and SDC1 protein confirmed by immunofluorescent staining (**Figure 4A** and **B**). We also confirmed the induction of CGB and SDC1 in transcript levels, both of which are well-known markers for ST subpopulations (Frendo et al., 2003; Jokimaa et al., 1998; Okae et al., 2018) (**Figure 4C**). More importantly, as ST secrete hCG into the maternal blood (Fournier, 2016), we verified by ELISA that our TSLC-derived ST secrete hCG significantly higher than TSLCs, reflecting the endocrinological capabilities of iPS-ST and ES-ST (**Figure 4D**). One of the main functions of placenta is acting as a barrier at the maternal-fetal interface, and ST is a major cell type composing that barrier. We evaluated whether TSLC-derived ST can function as a barrier by conducting the apical-to-basal leakage of sodium fluorescein (Na-Flu) assay (Rothbauer et al., 2017). iPS-TSLCs, ES-TSLCs and control TSCs were seeded in the apical chamber of transwell and cultured for 6 days in ST differentiation conditions. At days 3 and 6, the medium in the apical chamber was replaced with Na-Flu containing culture medium, and the cells were incubated for 1 hr. To assess the role of placental barrier in ST, we transferred the medium from the basal chamber to a 96-well plate and measured the levels of fluorescence. All tested cells showed similar reduction of the Na-Flu leakage at day 6, indicating that the ST derived from both iPS-TSLCs and ES-TSLCs can form a cellular barrier much like ST differentiated from genuine TSCs (**Figure 4E**). ST also plays an important role in active transport in the placenta by expressing various receptors and transporters (Cox, 2014; Tomi et al., 2011). In agreement with the previous reports, the expression of transport-related genes, such as ABCB1 and ABCG2 (well-known drug transporters in the placenta (Iqbal et al., 2012)), ABCC3 (ABC transporter in the placenta (St-Pierre et al., 2000)), and SLC15A1 (a solute carrier (Nishimura and Naito, 2005)) were elevated in both iPS- and ES-ST (**Figure 4F**). We further confirmed that ST differentiated from ES- and iPS-TSLCs showed increased expression of 303 ST-specific genes defined from first-trimester ST (Vento-Tormo et al., 2018) (**Figure 4G, Table S3**). Taken all together, these results demonstrate that the TSLCs generated from primed PSCs harbor bipotency, a core characteristic of TSCs.

**Figure 4.**
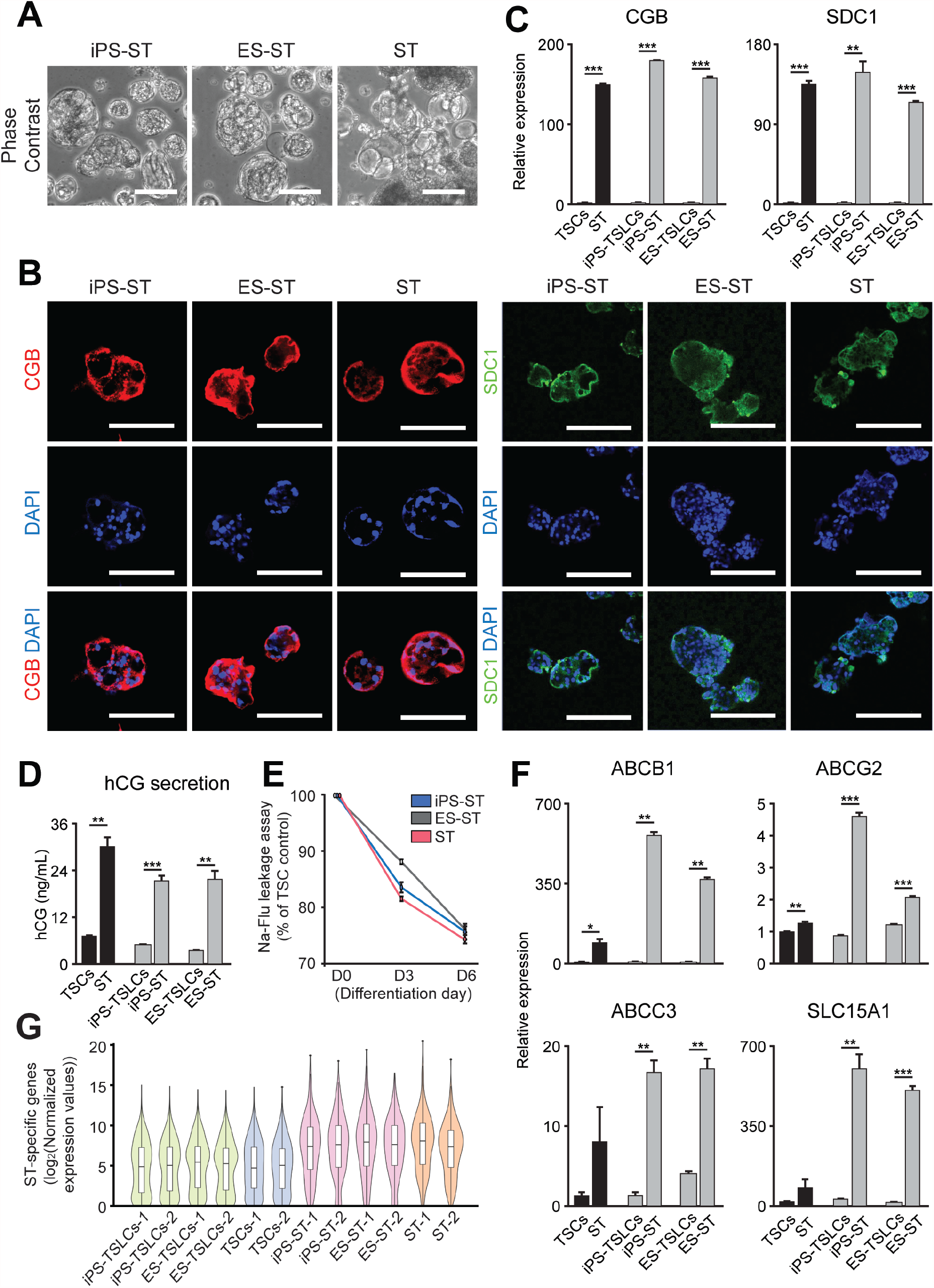
TSLCs can differentiate into ST-like cells. A) Representative phase contrast of indicated cell types. Scale bars indicate 100 µm. B) Representative IF images of indicated cell types. Red indicates the protein expression of CGB. Green indicates the protein expression of SDC1. Blue (DAPI) indicates the nuclei. Overlayed images are shown in bottom panels. Scale bars indicate 100 µm. C) Relative transcript levels of ST marker genes (CGB and SDC1) in indicated cell types to control TSCs. Error bars indicate SEM (n=3). Significance was indicated with **: p < 0.01 and ***: p < 0.001. D) The protein level of secreted hCG in indicated cell types. Error bars indicate SEM (n=3). Significance was indicated with **: p < 0.01 and ***: p < 0.001. E) Leakage of sodium fluorescein (Na-Flu) through iPS-ST, ES-ST, and ST differentiated on day 3 and 6 post seeding. Data are presented as mean ± SD for n = 2 and calculated as % of each day 0 control. F) Relative transcript levels of multiple ABC transporters (ABCB1, ABCG2, and ABCC3) and carrier transporter (SLC15A1) in indicated cell types to control TSCs. Error bars indicate SEM (n=3). Significance was indicated with *: p < 0.05, **: p < 0.01 and ***: p < 0.001. G) Violin plots showing the expression distribution of 303 first-trimester ST-specific genes (**Table S3**) (Vento-Tormo et al., 2018) in the indicated cell types. The box indicates 25th, 50th, and 75th percentiles.

## 4. DISCUSSION

Due to the difficulty and ethical issues of studying the human placenta *in vivo* as well as the unique aspects of human pregnancy compared to other mammals, it is imperative to develop a genetically tractable *in vitro* model of human placenta development (Myllynen and Vähäkangas, 2013). In this study, we successfully generated human TSLCs derived from human primed PSCs by optimizing the duration of BMP4 in TSCM. We verified that these TSLCs can self-renew for over a long period of time and have bipotency with which the cells can differentiate into EVT and ST. We have shown the global gene expression profiles of TSLCs, TSLC-derived EVT, and TSLC-derived ST were similar with those of human TSCs, EVT, and ST, respectively. We verified the expression of trophoblast-specific miRNA and hypomethylation of *ELF5* promoter in TSLCs. All these results indicate that these TSLCs accurately recapitulate two key features of TSCs, such as self-renewal and bipotency.

Multiple groups have also attempted to derive trophoblast lineage cells from human PSCs using BMP4 for the entire culture (Amita et al., 2013; Xu et al., 2002; Yabe et al., 2016). However, prior reports have mostly focused on the expression of differentiated cell markers, such as HLA-G or CGB, rather than those that typify stem cell properties. Although one group generated CT-like stem cells from human PSCs that express CDX2 and TP63 and confirmed that the resultant cells can be differentiated into ST-like and EVT-like cells based on the marker gene expression, invasion capacity, and hormone production, these cells do not efficiently self-renew and the authors did not test directed differentiation into ST or EVT (Horii et al., 2016). In accordance with this observation, we also confirmed that TSLCs generated by previously reported protocols undergo cell death after several passages, indicating they may not be suitable for long-term culture. In contrast, our TSLCs can be stably maintained without the loss of typical morphology of TSCs and key marker gene expression, indicating that we established a more selective and reliable method for deriving TSLCs from human primed PSCs.

Recently, two studies reported bipotent trophoblast-like stem cells derived from primed PSCs by micromesh and chemically defined culture medium, respectively (Li et al., 2019; Mischler et al., 2021). Additionally, there are multiple paths for deriving human TSCs from naïve ESCs (Cinkornpumin et al., 2020; Dong et al., 2020; Guo et al., 2021; Io et al., 2021) and reprogramming of somatic cells (Castel et al., 2020; Liu et al., 2020). Although these groups used different protocols for inducing trophoblast lineage cells from different cell sources, they showed that the induced cells exhibit typical characteristics of TSCs. Further systematic evaluation of the established cells and the culture protocols and will be critical for developing comprehensive *in vitro* placenta models.

Previous studies on inducing trophoblast lineage from human PSCs have resulted in different outcomes in some cases. Among others, two papers claimed that BMP4 treated PSCs are mesoderm cells expressing trophoblast marker genes (Bernardo et al., 2011) and amnion-like cells (Guo et al., 2021). The differences of protocols from previous papers are fibroblast growth factor 2 (FGF2) treatment with BMP4 and longer than 2 days of BMP4 treatment in different basal medium. Since we observed that the level of amnion marker gene was gradually increased upon longer BMP4 treatment (**Figure 1E**), we established our protocol by treating BMP4 for 2 days without FGF2 treatment. We demonstrated that our TSLCs have TSC-specific gene expression pattern comparable to TSCs rather than induced amnion-like gene expression patterns (**Supplemental figure 2B**), suggesting that our TSLCs formed cells with TSC-like properties, rather than amnion-like.

The ST plays important roles in metabolic activity of placenta by directly contacting with maternal blood, secreting hormones, and transporting nutrients for the fetus (Benirschke, 2012). Aberrancy in trophoblast lineage specification towards the ST may cause abnormal placental functions (Han et al., 2019). As one of the essential characteristics of ST is the expression of various drug transporters that may protect the fetus from xenobiotics, an appropriate *in vitro* placental model could be used for studying placental functions such as the regulation of drug transporters. Similar to the previous study (Yabe et al., 2016), we confirmed that TSLC-derived STs expressed multiple ABC transporters and carrier transporters, suggesting they can serve as a valuable model system to study placental transport and metabolic activity. In summary, we have demonstrated that self-renewing and bipotent human TSLCs can be derived from human primed ESCs and iPSCs. These TSLCs can serve as a useful *in vitro* model system to investigate the placental barrier and cell-based pharmacological screening for the transfer and biological effects of pathogens, drugs, or toxins.

## 5. CONCLUSIONS

Understanding the molecular mechanisms underlying human placenta development is crucial for developing prognostic and diagnostic measures as well as therapeutic interventions on various placenta-associated disorders. Until now, a limited access to a reliable human placenta model due to practical and ethical issues hampered our understanding of human placenta development. Here we successfully generated self-renewing and bipotent TSLCs from human PSCs using short-term treatment of BMP4 in TSCM. We propose that this protocol could be efficiently utilized to derive TSLCs from the pre-existing PSCs from patients with various conditions, thereby TSLCs can be used to genetically dissect the mechanisms of pregnancy complications during early development and will advance our understanding of the mechanisms underlying placenta development and placenta-related diseases.

## Author contributions

J.K. conceptualized the study. Y.J. and M.K. performed experiments. Y.J., M.K., and B.L. performed data analyses. Y.J., M.K., B.L., and J.K. wrote the manuscript.

## Acknowledgments

We would like to thank Dr. Takahiro Arima (Tohoku University) for sharing human TSC lines and Dr. Lucy LeBlanc for critical reading of the manuscript. This work was supported by R01GM112722 (NIH/NIGMS), R01HD101512 (NIH/NICHD), and Preterm Birth Research Grant (1017294) from the Burroughs Wellcome Fund to JK.

## FIGURE LEGENDS

**Supplemental Figure 1.**
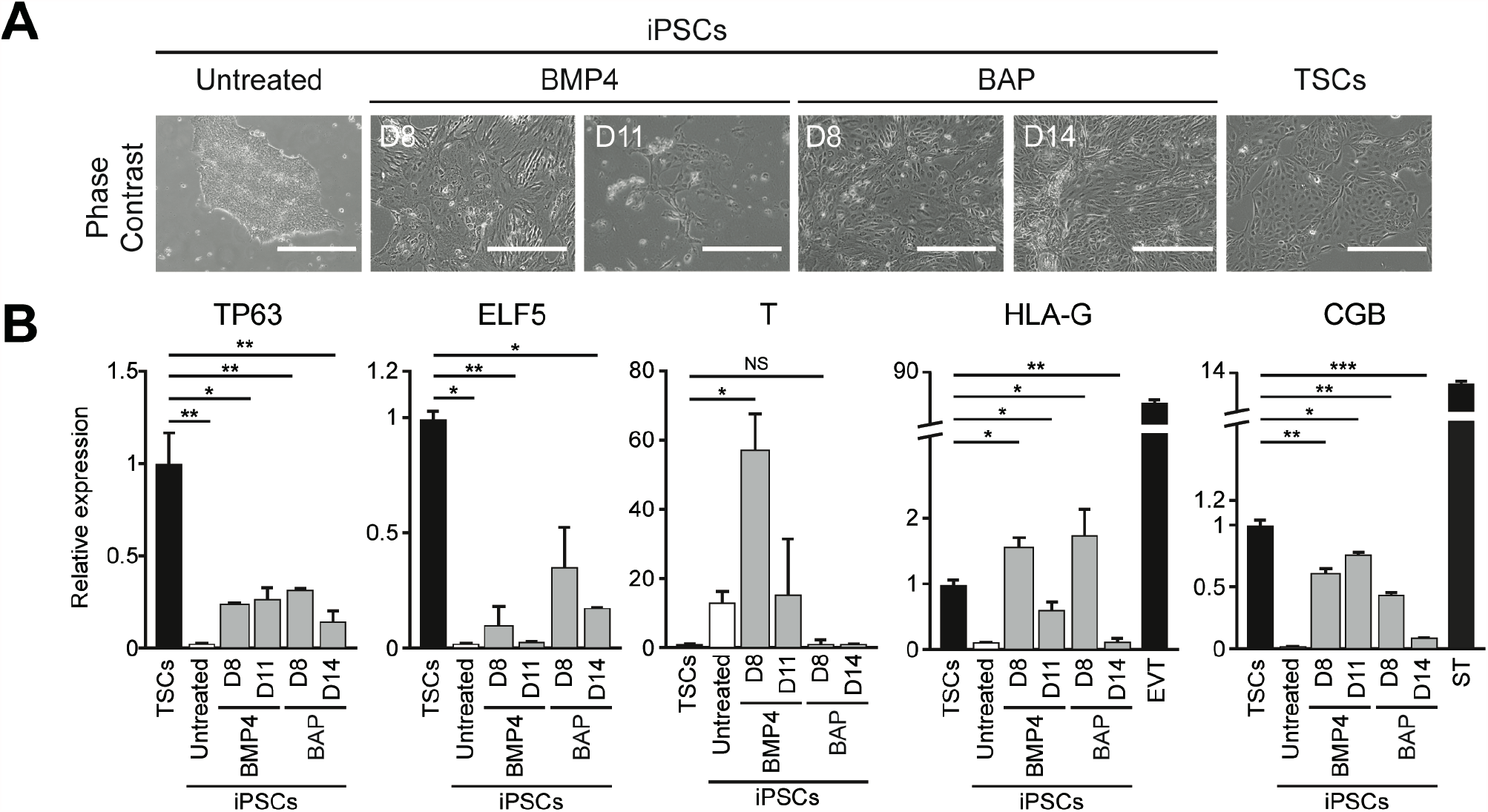
Continuous treatment of BMP4 or BAP. A) Representative phase contrast images of iPSCs (untreated), continuously BMP4- or BAP-treated iPSCs for up to 14 days, and TSCs. Scale bars indicate 400 µm. B) Relative transcript levels of TSC (TP63 and ELF5), mesendoderm (T), EVT (HLA-G), and ST (CGB) marker genes in untreated iPSCs and BMP4- or BAP-treated iPSCs for the indicated days compared to control TSCs. Error bars indicate SEM (n=3). Expression of HLA-G and CGB were also shown in EVT and ST, respectively. Significance was indicated with *: p < 0.05, **: p < 0.01 and ***: p < 0.001.

**Supplemental figure 2.**
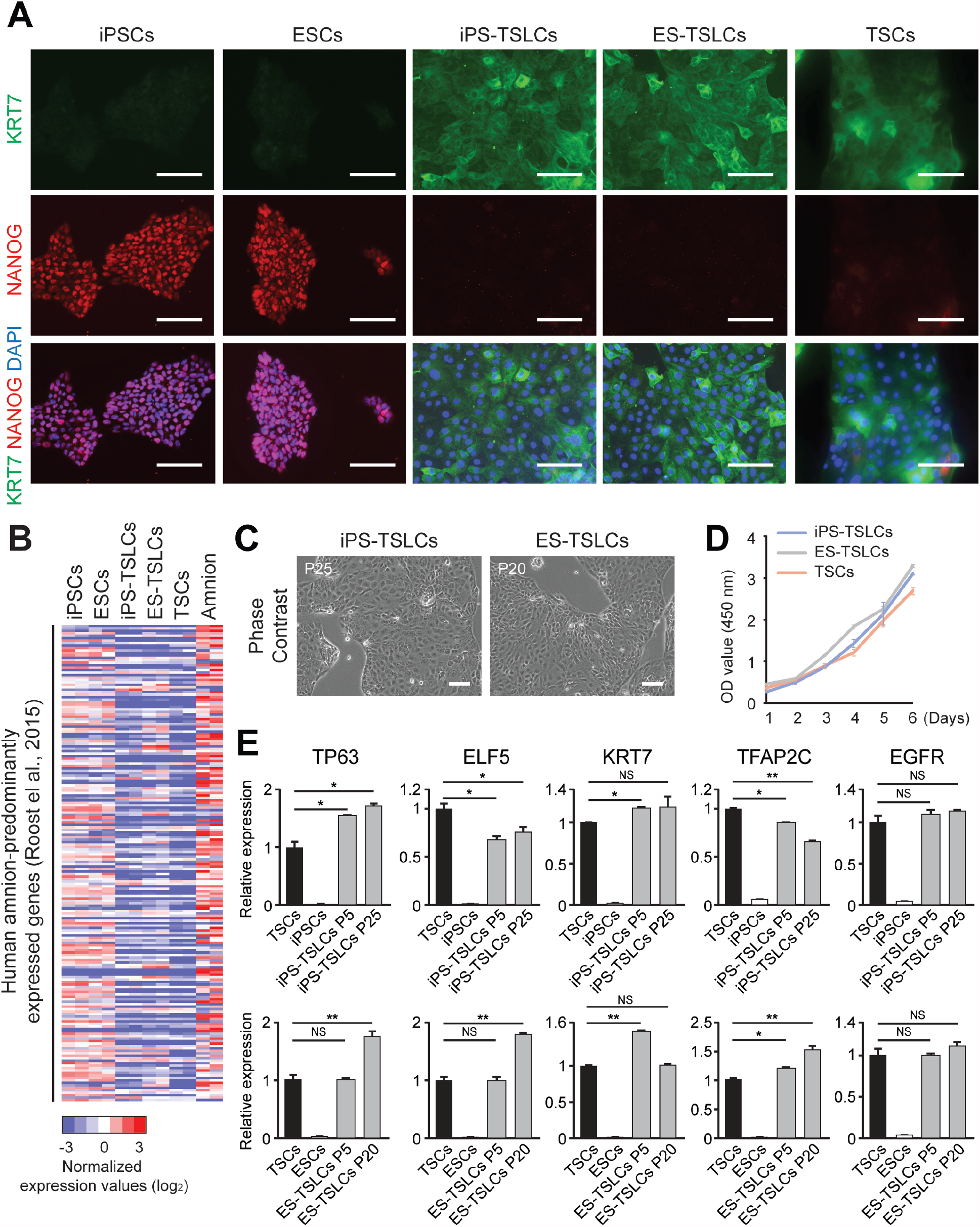
Self-renewal capacity of TSLCs after prolonged cell culture. A) Representative IF images of KRT7 and NANOG in PSCs, TSLCs, and TSCs. Green indicates the protein expression of KRT7. Red indicates the protein expression of NANOG. Blue (DAPI) indicates nuclei. Scale bars indicate 100 µm. B) A heatmap showing relative expression of 198 human amnion-specific genes predominantly expressed genes in amnion tissue at 9 gestational weeks (**Table S3**) (Roost et al., 2015) among iPSC, ESCs, iPS-TSLCs, ES-TSLCs, TSCs, and human amnion tissue (Roost et al., 2015). C) Representative phase contrast images of iPS-TSLCs and ES-TSLCs after over 20 passages, respectively. Scale bars indicate 100 µm. D) Line graphs showing the growth rate of the indicated cell types. E) Relative transcript levels of TSC markers (TP63, ELF5, KRT7, TFAP2C, and EGFR) in indicated cell types to control TSCs. Error bars indicate SEM (n=3). Significance was indicated with *: p < 0.05, **: p < 0.01 and ***: p < 0.001.

